# Quantifying temperature compensation of Bicoid gradients with a fast T-tunable microfluidic device

**DOI:** 10.1101/2020.03.09.983742

**Authors:** H. Zhu, Y. Cui, C. Luo, F. Liu

## Abstract

As a reaction-diffusion system strongly affected by temperature, the early fly embryos surprisingly show highly reproducible and accurate developmental patterns during embryogenesis under temperature perturbations. To reveal the underlying temperature compensation mechanism, it is important to overcome the challenge in quantitative imaging on fly embryos under temperature perturbations. Inspired by a microfluidics generating temperature steps on fly embryos, here we design a microfluidic device capable of ensuring the normal development of multiple fly embryos as well as achieving real-time temperature control and fast temperature jumps for quantitative live imaging with a home-built two-photon microscope. We apply this system to quantify the temperature compensation of the morphogen Bicoid (Bcd) gradient in fly embryos. The length constant of the exponential Bcd gradient reaches the maximum at 25 °C within the measured temperatures of 18-29 °C and gradually adapts to the corresponding value at new temperatures upon a fast temperature switch. Such an adaption decreases to a less degree if temperature is switched in a later developmental stage. This age-dependent temperature compensation could not be explained with the traditional synthesis-diffusion-degradation (SDD) model assuming the static parameters but an extended SDD model incorporating the dynamic change of the parameters controlling the formation of Bcd gradients.

**SIGNIFICANCE:** Thermal robustness is important for biological systems experiencing temperature fluctuations. To reveal the temperature compensation mechanism, the fruit fly embryo is an ideal model system. It is intriguing how the early fly embryo achieves highly reproducible and accurate patterning despite it is a reaction-diffusion system strongly affected by temperature. However, it has been challenging to quantitatively measure the developmental patterns in fly embryos under temperature perturbations. To overcome this problem, we construct a fast temperature tunable microfluidic device for fly embryos. Combining quantitative imaging with this device and mathematical modeling, we successfully quantify the temperature response of the morphogen Bicoid (Bcd) gradient and reveal that the temperature compensation for the Bcd gradient is stronger in the later developmental stage.

## INTRODUCTION

As a globe factor, temperature strongly influences activities of organisms especially for ectothermic animals without inherent temperature control. Interestingly, many important functions of the biological system usually show thermal robustness. For example, the circadian clock period is maintained to be constant under varied philological temperatures (1) and chemotaxis of *Escherichia coli* is robust against the ambient temperature variations (2). The thermal robustness of biological systems has been thought to be an intrinsic property of the gene regulatory network (3). To understand its origin, it is of importance to reveal the underlying temperature compensation mechanism.

One of the ideal model systems to study the temperature compensation mechanism is the *Drosophila melanogaster* (fruit fly) embryo (3–11). Since the fly is ectothermic, its embryos laid in the natural environment typically experience temperature fluctuations during at least one day/night cycle. Interestingly, the temporal development of fly embryos from different species scales uniformly (4, 6), and is highly robust across temperature (4, 6) The underlying mechanism to account for these phenomena is still under investigation. The temperature compensation mechanism in early embryos is even more intriguing. Since no cell membrane forms till the start of the 14^th^ nuclear cycle (nc14), i.e., 130 minutes after the embryo is deposited (AED) at 25 °C, the whole syncytial embryos can be treated as a reaction-diffusion system. This system is approximately an ellipsoid with the length of the principal axes of 500, 200 and 200 µm. As the temperature increases from 18 °C to 29 °C, the development time of embryos in the first 13 nuclear cycles shortens from 210 min to 110 min (4, 6), and the embryo laid by the flies raised in different temperatures for weeks also adapts to shorter length by 8% (12). One would expect that the developmental patterns in early fly embryos be much more strongly affected by temperature. However, the spatial developmental pattern along the anterior-posterior (AP) axis, e.g., the gap gene *hunchback (hb)* profiles, is robust against the temperature perturbations (11). Amazingly, one experiment shows that the pair rule gene profiles, e.g., Even-skipped strips, regularly form even if two halves of fly embryos are immersed at different temperatures with a microfluidic device generating a temperature step, i.e., sharp temperature gradient (9).

It is, however, still controversial what the temperature compensation mechanism is in the developmental pattern formation in fly embryos. Two hypotheses have been proposed. One suggests that the temperature compensation for the developmental patterns is originated from the cross-regulation of the downstream segmentation gene network. As the upstream regulator of *hb*, the maternal Bicoid (Bcd) gradients are observed to vary significantly at different temperatures (10, 11). And it has been suggested that a bi-gradient consisting of the Bcd gradient and an unknown posterior gradient might contribute to the temperature compensation (11, 13), but this hypothesis has been challenged (9), and the endo-siRNA pathway was discovered to be necessary for temperature compensation (13, 14). The other suggests that the adapted Bcd gradients could maintain the invariable Hb boundaries with the same activation threshold within a temperature range (8).

To clarify this debate, one of the keys is to solve the technical challenges in quantitative live imaging on patterning gene profiles under temperature perturbations. Ideally, one wishes to obtain the real-time dynamics of the Bcd gradient and its downstream gene profiles in response to the temperature perturbations. However, the developmental pattern of fly embryos dynamically evolves in minutes (15, 16) and the embryo is much larger than single cells, hence it has been difficult to uniformly change the temperature of the embryos in the time scale of seconds with most of the temperature control unit (17, 18). One promising approach would be the microfluidic device generating temperature steps (9, 10, 19). It enables the extraction of the nuclear fluorescence intensity of His2AvD-GFP or Bcd-GFP in live imaging on fly embryos (10). But it is still difficult to quantify Bcd gradients with a sufficient signal to noise ratio as demonstrated in previous work (20, 21). One reason could be the imaging plane inside the chip are several hundred micrometers below the microfluidic chip surface (19), beyond most of the work distance of the immersed objective of the microscope with a large numerical aperture (NA). Sophisticated microfluidic devices have also been developed to trap fly embryos for quantitative live imaging (22, 23), but it is still challenging to efficiently image the AP patterning of the fly embryo on the mid-sagittal plane.

Here we designed a new microfluidic device and established an imaging system based on a home-built two-photon microscope. With this system, we can uniformly heat or cool fly embryos in the temperature range of 18-29 °C, switch the temperature in a very short time, e.g., switch from 18 °C to 25 °C in 10 seconds, and run quantitative live imaging on the developmental patterns on fly embryos. We measured the Bcd gradients at the temperatures of 18 °C, 22 °C, 25 °C, or 29 °C, and analyzed the temperature dependence of the parameters of the Bcd gradient with the synthesis-diffusion-degradation (SDD) model (24, 25). We performed the fast temperature switch experiment and found that temperature compensation of fly embryos strengthens during early development. And this phenomenon could be explained only if we extend the SDD model by incorporating the time-dependence of the parameters controlling the formation of Bcd gradients. These results deepen our understanding of the temperature compensation mechanism of Bcd gradients. And this fast temperature-tunable (T-tunable) imaging system will facilitate investigating the temperature compensation mechanism of the pattern formation in fly embryos.

## MATERIALS AND METHODS

### Fabrication of the T-tunable microfluidic device

After the PDMS chip was cleaned with the tape (3M Scotch, Magic™ Tape, 810#) and the glass chip was purged with deionized water, they were put into the plasma cleaner (Harrick Plasma, Expanded Plasma Cleaner, PDC-001 (115V) | PDC-002 (230V)) simultaneously. To ensure the waterproofness of the assembled device, it is crucial to control the processing time of plasma cleaning as 90 ± 5 seconds in the low radio frequency level. After the glass chip was taking out for 30 seconds, the fly glue was deposited on the sample channel region with a pipette. Finally, the embryos were quickly stuck on the glass chip with the fly glue, and the microchip was assembled by pasting the PDMS chip and glass chip together.

The embryos laid by the fly line *egfp-bcd*;+;*bcd*^*E1*^ were collected for 1.5 hours at 25 °C with a fly cage as described before^(21, 26)^. To facilitate assembling, dechorionated embryos were mounted with the lateral orientation on a microarray under a stereomicroscope (Olympus, SZX7) before they were glued to a coverslip.

The temperature switch system consists of two water tanks (Echotherm™ IC20 Digital, Electeonic Chilling/heating Dry Bath), two peristaltic pumps (Masterflex L/S, Easy-Load II, Model 77200-62), two triple valves (Cole-Parmer, D062), connected by rubber hoses (Masterflex L/S 24) as shown in Figure 1A. The temperature was monitored with a thermocouple probe of digital thermometers (UNI-T UT 320) in the middle of the channel through a glue-sealed hole in the PDMS chip.

**FIGURE 1.**
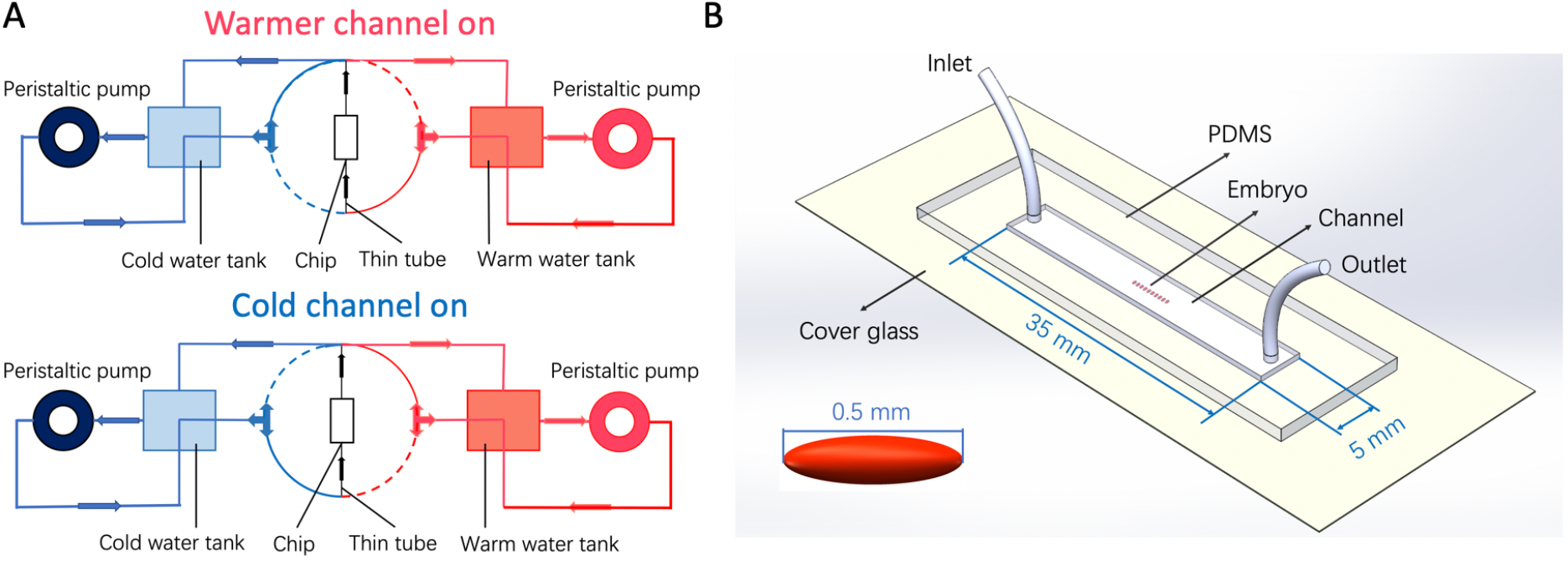
Schematic of the T-tunable microfluidics for live imaging on fly embryos. (A) Temperature-switching system. Two independent fluid paths feeding into the microfluidic chamber are switchable between the circulating loop (solid line) and the non-circulating loop (dotted line) by three-way valves. In each path, one peristaltic pump drives the flow with a specific temperature controlled by a water tank. (B) Microfluidic chip for live imaging on fly embryos. Multiple fly embryos (~0.5×0.2×0.2 mm^3^) are glued on a cover glass (0.13~0.16 mm in thickness, CITOGLAS, 80340-6910) and immersed in a microfluidic chamber (L*W*H=35*5*0.6 mm^3^) made of PDMS.

### Simulation on microfluidics

The fluid flow and temperature distribution in the microfluidic channel were simulated with the software COMSOL (COMSOL Multiphysics 5.2a). The three-dimensional geometry of the whole model was nearly identical to the device as shown in Figure 1B. The material of the upper and bottom of the chip was set as PDMS and glass, respectively. The embryo was simplified as an ellipsoid full of liquid water surrounded by a wax layer with a thickness of 0.3 µm and the thermal conductivity *k* = 2.5×10^−3^ kJ/(m· s · K), which represents the waxy layer and the vitelline membrane of dechorionated embryos (26). The other part of the channel was full of liquid water and the flow rate was set based on the experimental value, e.g., 3.3 cm/s corresponding to ~6 mL/min. The entire system was set as the closed no-slip boundary condition except for the outlet and inlet. And the whole model was set as the physics-controlled mesh of sequence type with a finer element size. The temperature was set as 18 °C, 22 °C, 25 °C, or 29 °C in the inlet, and the temperature surrounding outside of the chip as the room temperature of 20 °C. All models here assumed the system was operating in a steady state.

### Imaging experiment

The water tanks were switched on and set as the default temperature in advance. The plastic tubes were connected with the chip’s inlet and outlet, and ensure there was no leaking water. Then, the temperature control system was assembled as displayed (Fig. 1A). Embryos were imaged with a home-built two-photon microscope (27). Microscope control was implemented using ScanImage 3.8 (28). The images were captured with a Zeiss 25× (NA 0.8) oil/water-immersion objective and a gallium-arsenide-phosphide (GaAsP) photomultiplier tube selected for the background less than 4000 counts/s. The excitation wavelength was 920 nm and the average laser power at the specimen was 28 mW. Each complete embryo image was stitched with three images (512×512 pixels, with 16 bits and at 2 ms per line) along with the AP axis of the embryos, and each image was an average of three sequentially acquired frames to improve the signal to noise ratio. The Bcd-GFP gradient was imaged in the mid-sagittal plane at 28 min, 21 min, 15 min, and 11 min into nc14 as the measurement temperature was 18 °C, 22 °C, 25 °C, or 29 °C, respectively. Since the measurement temperature could be different from the embryo collection temperature, the embryos were chosen only if they were younger than nc11 at the beginning of the imaging session to ensure sufficient adaption time in the setting temperatures before measurements.

### Image analysis

Images are processed and analyzed with customized MATLAB codes (MATLAB 2018a). The nuclei band of the embryo was identified with a classical morphology algorithm. The interval of each nucleus and the center positions were determined by transforming the nuclei intensity into the frequency domain with Fast Fourier Transform (FFT). The long axis and short axis of each nucleus were identified with shape recognition and the average fluorescence intensity was extracted. The fluorescence background was nearly zero based on the measurement of the wild type embryos (*w^1118^*) without Bcd-GFP expression under the same imaging condition.

### Modeling

As the GFP protein needs tens of minutes to mature to fluorescence, the fluorescent Bcd-GFP gradient profile observed in live imaging deviates from the total Bcd-GFP gradient profile (20, 29, 30). The dynamics of the gradient can be depicted with the SDD model incorporating the maturation correction (20). At the steady state, the concentration of immature proteins with no fluorescence *c*_*im*_(*x*) = *c*_0_ * *k* * *e*^−*x*/(*k*\**λ*)^, where 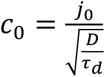, 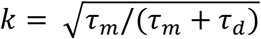, where *j*_0_ is the Bcd-GFP synthesis rate and *τ*_*m*_ was the mature time of Bcd-GFP. And the observed Bcd-GFP gradient in live imaging is contributed by mature proteins with the concentration *c*_*m*_(*x*) = *c*_*tot*_(*x*) − *c*_*im*_(*x*), where *c*_*tot*_(*x*) represents the total Bcd-GFP. Since the temperature varied in a small range in the measurement, the diffusion constant was determined with a simplified formula: 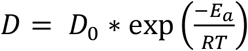, given the measured diffusion constant *D*= 3.60 µm^2^/s at 18 °C (29) and assume *E*_*a*_ = 1.67×10^4^J/mol (31), *D*_0_=3.66×10^3^µm^2^/s. On the one hand, the degradation time of Bcd-GFP at different temperatures could be calculated based on 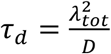, where *λ*_*tot*_ is the maturation corrected length constant of the Bcd-GFP gradient. On the other hand, based on the SDD model incorporating with maturation correction, *λ*_*tot*_ needs be derived from the Bcd-GFP gradient measured with live imaging given the degradation time and the maturation time of GFP. To reduce the flexibility, the GFP maturation time is also assumed to be consistent with the Arrhenius equation, i.e., 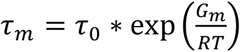, where *τ*_0_ is a constant and *G*_*m*_ is the activation energy for maturation. Based on the previous measurements, we estimate the maturation time at different temperatures by taking *τ*_*m*_ = 50 min at 22°C (32, 33) and *G*_*m*_=4×10^4^J/mol, which is consistent with the temperature dependence of the maturation time of mEGFP in *E.coli* (34) (mEGFP has one extra mutation to suppress the dimerization of eGFP, and their fluorescence properties are nearly the same (35)). Self-consistent *λ*_*tot*_ and *τ*_*d*_ can be obtained with iteration according to the flow chart (Fig. S3). To calculate 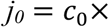 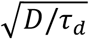 at different temperatures (Table 1), the fitted *C_0_* with an arbitrary unit was converted to *c*_0_ with the unit of the absolute molecule number per unit volume. Based on the previous live imaging experiment, the maximum absolute concentration of nuclear Bcd-GFP in *egfp-bcd*;+;*bcd*^*E1*^ fly embryos was 36 molecules/µm^3^ at room temperature (21) (taking it as 22 ℃). And the maximum concentration of the Bcd-GFP gradient is at ~10% EL (30). Based on the maturation correction model, we acquired the corrected gradient amplitude 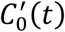 at different temperature *T*. Hence 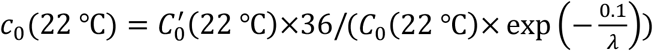 and 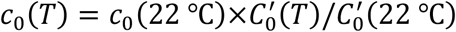.

**TABLE 1.** Temperature-dependent parameters of the traditional SDD model. *T* represents the measurement temperature. *λ* and *C_0_* represent the length constant and amplitude of the Bcd gradient from exponential fits in the region 50 µm to 250 µm away from the anterior pole, respectively. *λ*_*tot*_ represents the maturation corrected length constant. *D*, *τ*_*m*_, *τ*_*d*_ and *j*_0_ denote the diffusion constant, maturation time, degradation time, and the synthesis rate of Bcd-GFP at the anterior pole calculated with the absolute molecule number after maturation correction (20, 21), respectively (see Materials and Methods).

| $T$ (K) | $\lambda$ ($\mu\text{m}$) | $\lambda_{tot}$ ($\mu\text{m}$) | $\ln(C_0)$ | $D$ ($\mu\text{m}^2/\text{s}$) | $\tau_m$ (min) | $\tau_d$ (min) | $j_0$ (molecules /$\mu\text{m}^2 \cdot \text{s}$) |
| --- | --- | --- | --- | --- | --- | --- | --- |
| 291 | 70.7 | 58.7 | 7.4 | 3.60 | 62.2 | 16 | 20.5 |
| 295 | 80.3 | 65.9 | 7.3 | 3.95 | 50 | 18.3 | 14.7 |
| 298 | 96.4 | 80 | 6.8 | 4.23 | 42.1 | 25.2 | 7 |
| 302 | 77.2 | 64.8 | 7.3 | 4.63 | 34 | 15.1 | 14.5 |

The synthesis rate, diffusion constant and degradation rate of Bcd proteins dynamically change during embryogenesis, hence the SDD model is extended to incorporate these time-dependent parameters (Fig. S6). The maximum of these parameters and the maturation rates are kept the same as the traditional SDD model (Table S1). And the spatio-temporal evolution of Bcd gradients is obtained via numerical simulations. Through systematic search based on the published experimental results (29, 30, 33), a set of optimized parameters can be identified to match the simulated Bcd profiles well with the dynamics of normalized Bcd-GFP profiles measured with fixed embryos and the Bcd-GFP profiles measured with living embryos at 22 ℃, and the normalized Bcd-GFP gradient measured with living embryos at the equivalent time (corresponding to 15 min at 25 ℃) into nc14 at four temperatures. And the extended SDD model is consistent with the length constant changes upon the temperature switches at early and late developmental time. In the parameter sensitivity analysis, each parameter is increased or decreased by 0-90% or 0-30% of its optimized value, the goodness of fit is calculated with either the mean square error (MSE): 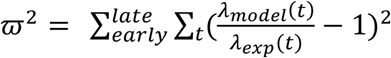 or the cross entropy loss 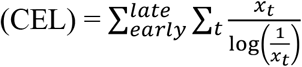, 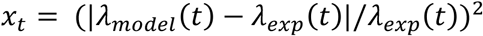.

## RESULTS

### Device design

Given the problems in the past experiments, we hope to achieve the following goals in designing this microfluidic chip: 1) live imaging: the chip has good gas permeability and the water flow is large enough to deliver sufficient oxygen to ensure the normal development. On the other hand, the flow rate of water cannot be too large, otherwise, the severe hydrodynamic pressure will also affect the development of the embryo. Finally, the imaging plane of the embryo is within the working distance of the objective of the microscope. 2) real-time temperature control: temperature inside the embryo is uniformly distributed and can be switched in tens of seconds.

To achieve the above two goals, we designed a microfluidic chip shown in Figure 1. The upper part of the chip is made of polydimethylsiloxane (PDMS), a transparent elastomer widely employed in biological microfluidics. According to the embryo size (~0.5 × 0.2 × 0.2 mm), the sample chamber is a rectangular channel with a width of 5 mm, a length of 35 mm and a height of about 0.6 mm. And the lower part of the chip is a microscope cover slide, on which multiple embryos are stuck with fly glue and PDMS is fixed with plasma treatment to form a sample chamber without any water leakage. Hence the image plane at the middle of the embryo is within the working distance (0.4 mm) of the objective.

To control the developmental temperature of embryos, the embryos inside the microfluidic sample chamber are immersed in flowing water. The inlet and outlet plastic tubes of one fluid loop are connected with two circular openings with an outer diameter of 1.3 mm in PDMS (Fig. 1B). To ensure the rapid circulation of water, inside the loop one peristaltic pump drives the flow through the tube. And water can be heated or cooled by a water tank. To minimize the heat dissipation during circulation, the fluid path is as short as possible. To switch the temperature, two independent fluid paths with different temperatures can be switched with three-way valves to feed into the microfluidic sample chamber to control the developmental temperature of embryos (Fig. 1A).

We choose to set the flow rate as 6 mL/min for the following reasons. Firstly, based on the Reynolds number (Re) Re = ρvd/µ, where ρ, v, d, and µ are the liquid density, flow rate, channel diameter, and viscosity coefficient, respectively, the flow in the channel is considered as a laminar flow as the Reynolds number is only 118, far less than 2000. This is confirmed in COMSOL simulations (Fig. 2A). Secondly, the calculated shear stress is only 0.33 N/m^2^, well below the damage threshold on embryos (36). And we found that inside the microfluidic chamber with such a flow rate the embryo development is indeed normal (Fig. S1) and the average hatching rate is 73± 5%. Thirdly, we confirmed that this flow rate ensures the stability of the temperature inside the chip. Based on the COMSOL simulations, the temperature of the fluid inside the sample chamber is uniform, i.e., the difference in the vicinity of the embryo is less than 0.1 °C (Fig. 2B). The temperature of the fluid inside the chamber slightly offsets from the one in the water tank and the difference drops as the flow rate increases. This temperature difference is caused by the heat dissipation from the water flow to the environment in the tubing and the sample chamber. The heat dissipation is greater if the objective is in contact with the cover glass, hence the temperature difference increases by 1 °C at the flow rate of 6 mL/min. Lastly, we also measured the response curve of the water inside the sample chamber upon temperature switches. As shown in Figure 2C, the response time for the jump from 18°C to 25 °C increased from 30 seconds to 10 seconds when the flow rate is increased from 2 mL/min to above 6 mL/min. And the heat transport between the embryos and the surrounding water reaches equilibrium in less than 0.1 s according to the COMSOL simulations. So our device can achieve both fast temperature-tunable control and live imaging for fly embryos.

**FIGURE 2.**
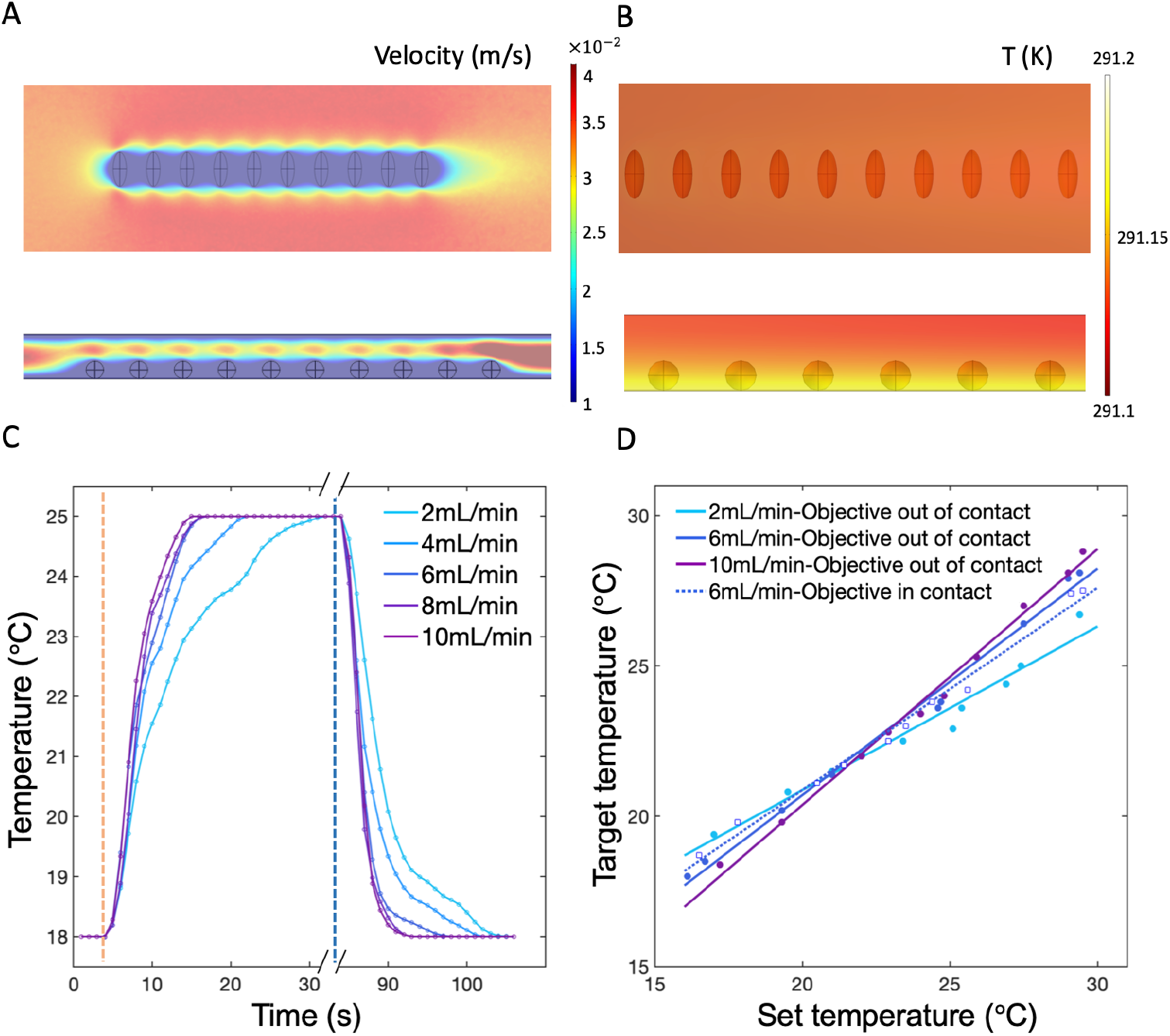
Characterization of the fast T-tunable microfluidic device. (A-B) Top view (up) and side view (down) of the simulated flow rate distribution (A) and temperature distribution profiles (B). (C) Temperature switching time series with different flow rates. The response time upon jump-up (left) and jump-down (right) between 18 °C and 25 °C is around 10 s when the flow rate is larger than 6 mL/min. (D) The target temperature measured in the chip as a function of the set temperature measured in the water tank under different flow rates with the microfluidic chip in contact with (circles as the mean values of the measurement results, solid lines as the linear fitting to the data) or without (dotted line, square) the objective lens at ambient temperature (20 °C).

### Temperature characteristic of Bcd gradients

Using this new microfluidic device, we detected the Bcd-GFP gradient profiles in fly embryos with a home-built two-photon microscope at temperatures of 18 °C, 22 °C, 25 °C, or 29 °C (see Materials and Methods), and we extracted the nuclear Bcd gradient profiles along the AP axis on the dorsal side of embryos (Fig. 3A). Bcd forms an exponential gradient from the anterior to the posterior pole. Based on the traditional SDD model (24, 25), the steady-state Bcd gradient shows a length constant 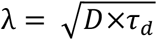, where *D* and *τ*_*d*_ are the diffusion constant and degradation time of Bcd proteins, respectively, which are both temperature dependent (see Materials and Methods). So we calculated the length constants by fitting the Bcd gradient profiles to an exponential function (Fig. 3B). Results showed that the average length constants of the Bcd gradient of individual embryos are 70.7 ± 10.4 µm at 18 °C, 80.3 ± 8.3 µm at 22 °C, 96.4 ± 21.4 µm at 25 °C, and 77.2 ± 9.1 µm at 29 °C. The length constants at 18 °C, 22 °C and 29 °C are significantly different from the one at 25 °C (*p* valves < 0.001). These results show that the length constants of Bcd gradient profiles are indeed temperature sensitive.

**FIGURE 3.**
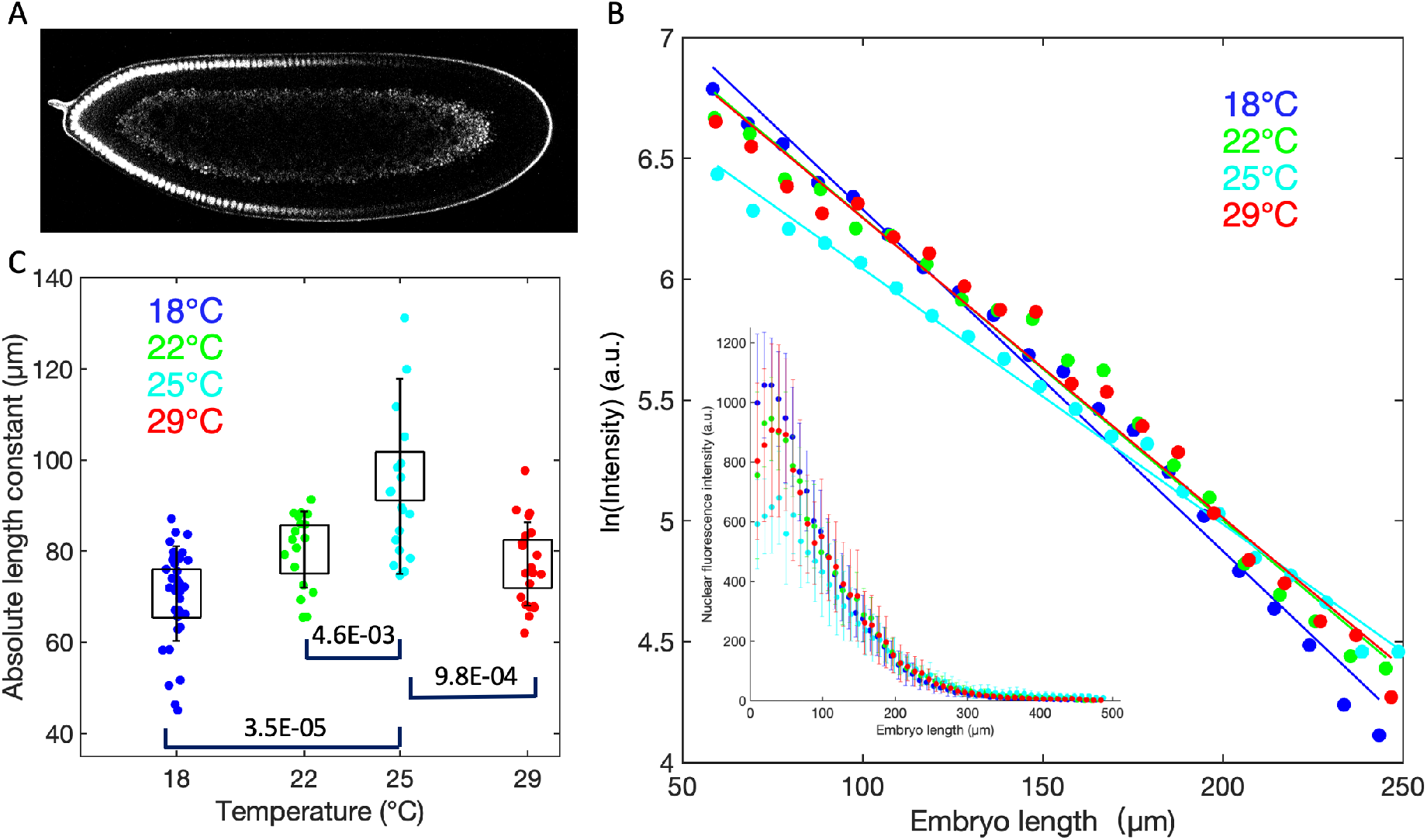
Temperature dependent Bcd gradient profiles. (A) A representative image of a fly embryo expressing Bcd-GFP proteins in the microfluidic chip at 22 °C measured with a home-built scanning two-photon microscope. (B) Logarithm of the average fluorescence intensity (background subtracted) of the nuclear Bcd-GFP in the embryo’s dorsal side as a function of the absolute embryo length (the anterior pole as 0) at different temperatures. Inset shows the whole range of background-subtracted nuclear Bcd-GFP intensity profiles. (C) The length constants of the Bcd gradient of individual embryos (colored circles) vary at different temperatures (blue, sample number *n* = 37 at 18 °C; green, *n*=18 at 22 °C; cyan, *n*=20 at 25 °C; and red, *n*=22 at 29 °C). Square boxes and error bars represent the mean values and the standard deviation, respectively. And the *p*-values are also shown for comparisons with the data at 25 °C.

Interestingly, the length constant increases to reach the maximum value at 25 °C then decreases at 29 °C. These results are consistent with previous immunostaining results in fixed embryos (8) as shown in Figure S2. Notably, both results show the same non-monotonic trend either on absolute length constants or the relative length constants (normalized by the embryo length). The slight discrepancy between the two measurement results could result from two factors. One is that the length constant of the Bcd gradient is not scaled with the fluctuating embryo length (11, 37). The other is that the Bcd gradient measured in live imaging still needs maturation correction for eGFP and the maturation time of eGFP could vary at different temperatures (38). However, one would expect that the length constant positively correlates with temperature assume both the diffusion constant and the degradation rate increase with the rise of temperature. It is well known that the diffusion constant of molecules in solutions follows the Stock-Einstein equation (39) and the solution viscosity follows the Arrhenius equation

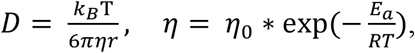

where *k*_*B*_ and *R* are the Boltzmann constant and the gas constant, respectively, and *η* is the solvent viscosity, which can be determined by the activation energy *E*_*a*_. We calculated the diffusion constant and the degradation time of Bcd-GFP at different temperatures and listed the results in Table 1 (for more details, see Materials and Methods). The calculated degradation time of Bcd-GFP is consistent with previous measurement results (21, 29, 33, 40). Interestingly, our results suggest that the degradation time does not follow the Arrhenius equation and it is maximal at 25 °C, which is the most suitable temperature for fly growth. To confirm the above predictions, it worth investigating the temperature dependence of these parameters in the future.

### Perturbation of the Bcd gradient upon temperature switches

With our newly developed microfluidic device, we can measure whether the Bcd gradient could dynamically adjust upon temperature perturbations during embryo development. Since the length constant difference is maximal between 25 °C and 18 °C, we switch the sample chamber temperature to 18 °C after the embryos have been developing at 25 °C for at least one hour. Surprisingly, as shown in Figure 4, if the switch time is earlier, e.g., 110 min AED, the length constant increases slightly but fails to reach the corresponding value at 25 °C (the temperature before the switch), then drops to the corresponding value at the new temperature of 18 °C. In contrast, when switching later, e.g., at 130 min AED, the length constant remains at the corresponding value at 25 °C after the temperature switches to 18 °C. These results indicate that the length constant of Bcd gradients adapts to a less degree if the temperature switches at a later developmental time, suggesting the Bcd gradient is more robust against temperature perturbations in the later developmental stage.

**FIGURE 4.**
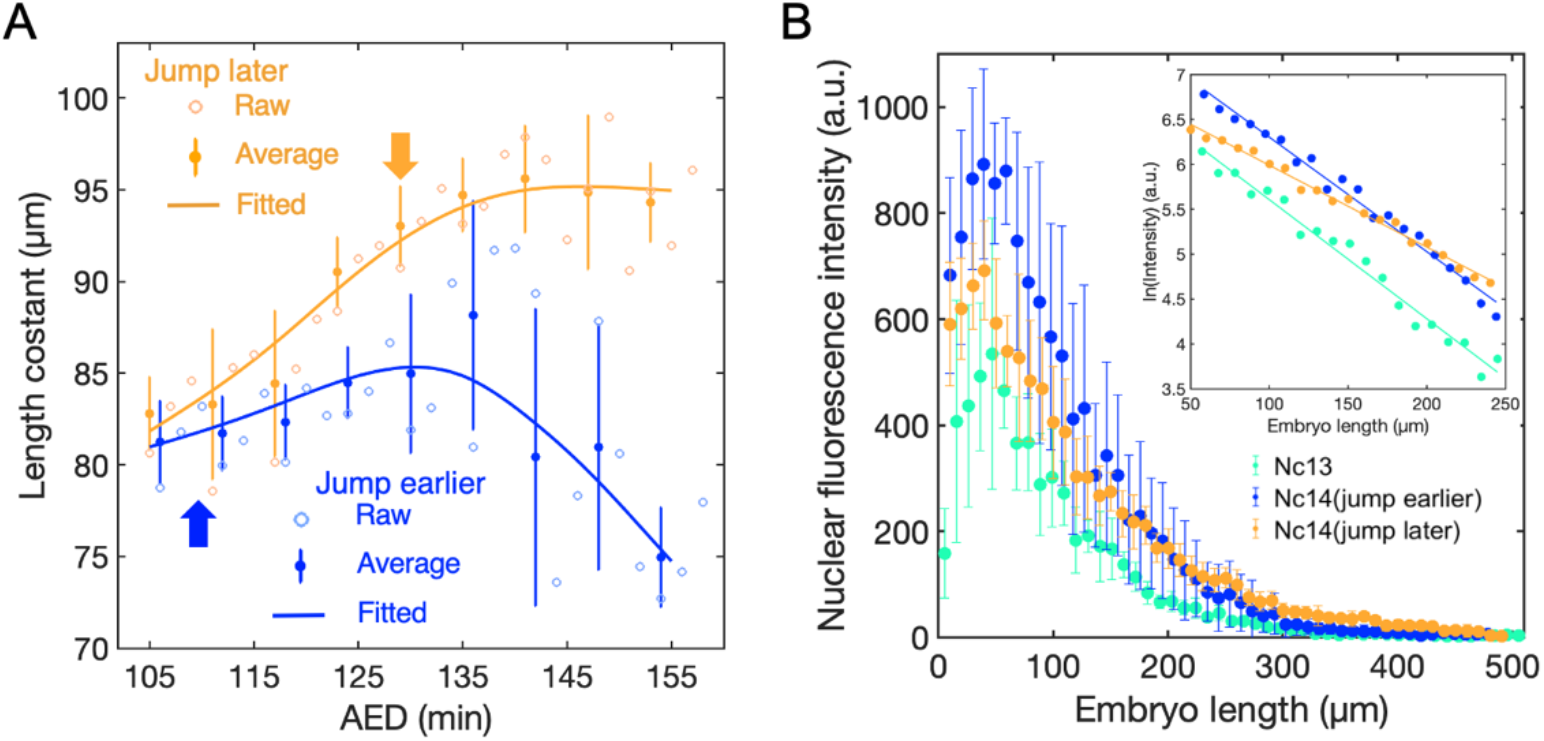
Response of Bcd gradients upon fast temperature switches. (A) The change of the length constants of Bcd gradients upon a temperature switch from 25 °C to 18 °C at 130 min AED (orange, arrow) and 110 min AED (blue, arrow), respectively. The dots represent the average of raw data (open circle) and the error bars represent the standard deviation (sample number *n*=3). The solid lines are the smoothing curve of the dots for eye guides. Notably, the development time at 18 °C is rescaled to match with the one at 25 °C for a better comparison of the temperature switches at different times. (B) The Bcd gradients (background subtracted) at about 6~12 min into nc13 (cyan), at around the 15 min into nc14 when jumping at 110 min AED (blue) or 130 min AED (orange). Error bars represent the standard deviation. The inset shows the logarithm of Bcd gradients (background subtracted).

To understand the above observation in live imaging, we used the traditional SDD model incorporating maturation correction to calculate the dynamic change of the Bcd gradient upon the temperature switch at different developmental time points (see Materials and Method and Fig. S3). We simulated the dynamics of the Bcd profiles at different temperatures. The simulated Bcd gradient profiles seem to be consistent with the measured ones (Fig. 5A). For comparison, we rescaled the development time at different temperatures (Tables S1 and S2) and obtained the dynamic of the length constant of Bcd-GFP gradient in live embryos (Fig. 5B).

**FIGURE 5.**
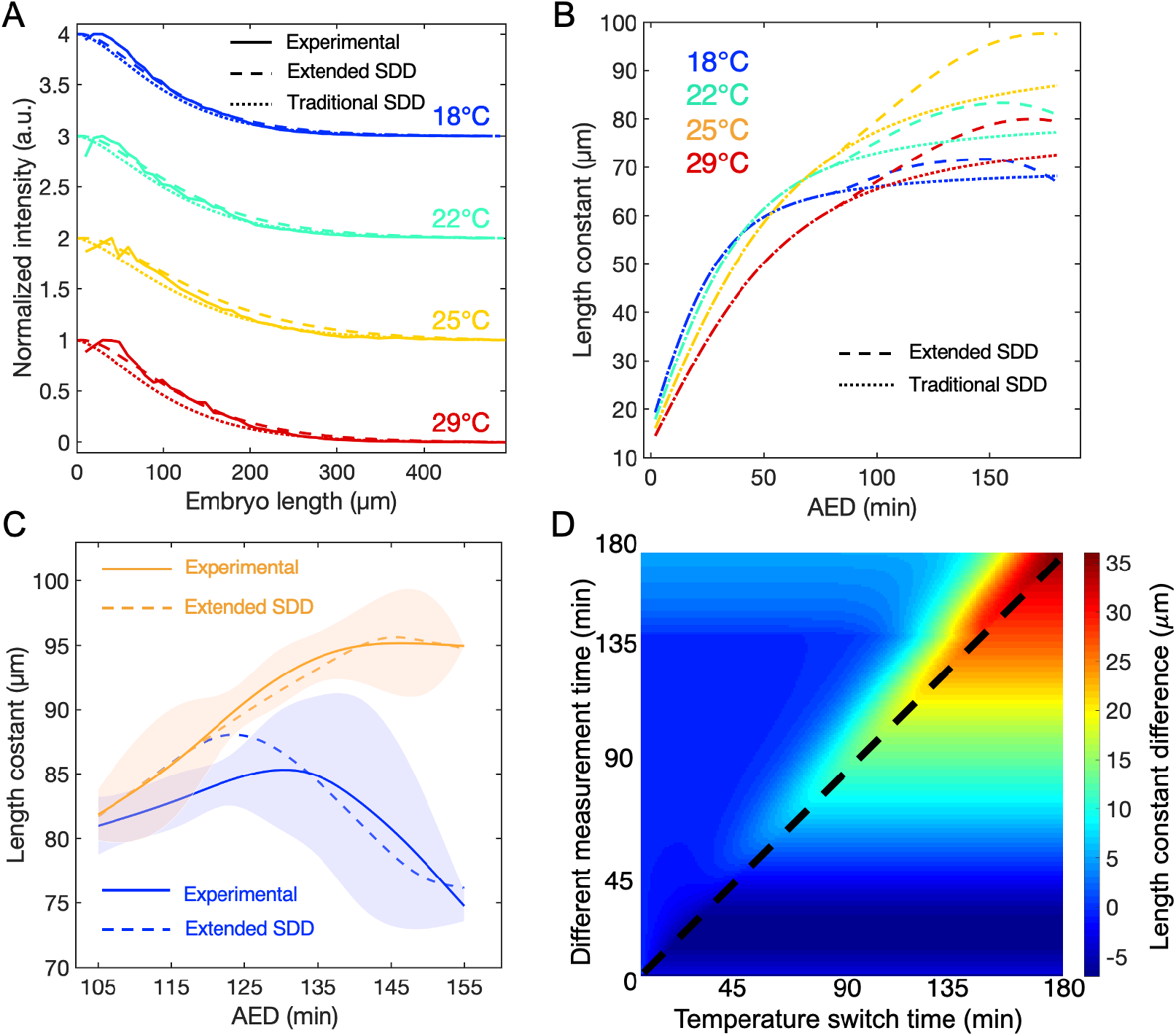
Simulations of SDD models upon temperature switch. (A) The normalized simulated Bcd gradients based on the extended SDD model (dashed line) fits better than the one based on the traditional SDD model (dotted line) to the normalized measured average Bcd gradients (solid line) at 26 min, 20 min, 15 min, and 11 min into nc14 at 18 °C (blue), 22 °C (cyan), 25 °C (yellow) and 29 °C (red), respectively. The measurement times match with each other after rescaling the developmental time at different temperatures. Profiles for different temperatures are shown with 1 offset on the normalized intensity axis to enhance visual clarity. (B) The fitted length constant of the maturation corrected live Bcd-GFP gradients based on the simulation of the extended (dashed line) or traditional (dotted line) SDD model as a function of the time after embryo deposition (AED) at 18 °C (blue), 22 °C (cyan), 25 °C (yellow), and 29 °C (red). (C) Simulated length constants based on the extended SDD model (dashed line) agree well with the experimental length constants (solid line) as the temperature switches earlier (blue) or later (orange). The simulated length constants after temperature switch are averaged by varying the switch time ±1 min considering the measurement error of the switch time. The shade represents the standard deviation of the measured length constants as shown in Fig. 4A. (D) Heatmap (colormap unit: µm) for the simulated length constant differences for the Bcd gradient at the initial temperature of 25 °C with the respect to that of 18 °C before (below the dashed line) and after (above the dashed line) the temperature is switched to 18 °C.

However, the simulated length constants change upon temperature switches cannot fit the experimental results after an exhaust search, and the adaptation in response to a late temperature switch is the same as the early ones (Fig. S5A). This discrepancy might be attributed to the time dependence of the key parameters in the SDD model. First, it has been shown that the synthesis rate of the Bcd increases after embryo deposition and decreases significantly in early nc14 due to the degradation of the *bcd* mRNA (30, 40) secondly, it was also suggested that the degradation time of Bcd changed before and after nc14 (33, 40); finally, the diffusion of Bcd could also slow down, as the Bcd molecules are pumped into the nuclei (41) and the nuclei number keeps increasing after each cycle, and more importantly, the formation of the cell membrane in nc14 could constrain the transport of Bcd. Most of these phenomena are related to the maternal to zygotic transition (42), which happens at around the start of nc14.

To further explore the underlying mechanism, we exploit an extended SDD model incorporating the dynamics of the key parameters controlling the formation of Bcd gradients (Fig. S6). Based on this extended SDD model, the simulated Bcd profiles (Movie S1) can match the dynamics of measured Bcd profiles with fixed embryos after intensity normalization (Fig. S7), the Bcd-GFP profiles measured with live imaging fit better than those based on the traditional SDD model with our measurement results at four temperatures after intensity normalization (Fig. 5A). More importantly, the dynamics of the simulated length constants of Bcd-GFP gradients upon the temperature switches at early and late developmental time (Movie S2 and S3) are consistent with the experimental results (Fig. 5C). Using this set of parameters (Tables S3), we calculated the heat map showing the length constant difference upon temperature switches at other different developmental times (Fig. 5D). It shows that the length constant difference between two developmental temperatures becomes more significant as the developmental time increases. Moreover, it predicts the difference will diminish to a less degree if the temperature changes from 25 °C to 18 °C at a later developmental time. In contrast, the heat map calculated based on the traditional SDD model does not show the strong dependence on the switch time (Fig. S5B). Hence the extended SDD model can account for the strengthening of the temperature compensation effect on Bcd gradients at later developmental stages. Although the fitting to the temperature switch experiments might be further improved if we try a more extensive parameter search, the parameter sensitivity analysis indicates the fitting with the current set of parameters, which is consistent with the known experimental results (29, 30, 33), reaches a minimum (Fig. S8). Moreover, the switch time of the degradation time is the most sensitive parameter, followed by the switch time of the synthesis rate and the diffusion constant of Bcd. Whereas the change rates of these three parameters are much less insensitive. These predictions could help to design temperature-dependent dynamic measurements on these parameters, which could facilitate refining the extended SDD model to explore the temperature compensation of Bcd gradients in future studies.

## DISCUSSION

To understand the robustness of the regulatory network, it is important to investigate how organisms achieve reproducible patterning by temperature compensation under temperature perturbations. Our new fast T-tunable microfluidic device overcomes the limitation of previous devices (9, 19) and can realize better live quantitative imaging on fly embryos at different temperatures as well as fast temperature jumps. With this device, we measure the Bcd gradients of fly embryos at different temperatures, and compare the temperature dependence of the parameters of the Bcd gradient with the SDD model (24, 25). According to quantitative measurement results, the length constant of the Bcd gradients reaches the maximum at 25 °C. Upon the temperature jump in early developmental stages, the length constant of the Bcd-gradient gradually shifts from the value of the prior temperature to that of the new temperature. However, if the jump time is close to nc14, the length constant of the Bcd gradient could remain the same as the one before temperature jumps, suggesting that the temperature compensation for patterning strengthens during early fly embryogenesis.

Although the traditional SDD model assuming static parameters could account for the temperature-dependence of the Bcd gradient at specific development times, its prediction on the length constant changes upon temperature jumps is inconsistent with the experimental results. To account for the age-dependent temperature compensation effect on Bcd gradients, it is necessary to extend the SDD model by incorporating the time-dependent parameters. It would be interesting to test this prediction by quantifying the potential time-dependence of the synthesis rate, diffusion constant, degradation time, and maturation time of Bcd-GFP at different temperatures. This could deepen understanding of the dynamic formation and temperature compensation of Bcd gradients. Resonating with the recent experiments (20, 29, 43, 44) and modeling (45), it also suggests it is very important to investigate the dynamics of the Bd system in its formation and interpretation.

Furthermore, besides the Bcd gradient, it is also important to quantify the downstream patterning genes with the MCP-MS2 (46, 47) or LlamaTags (16) system in live imaging without maturation delay. Most importantly, we will be able to quantify the response of the patterning system in fly embryos with live imaging by applying a variety of dynamical temperature perturbations, e.g., periodic temperature fluctuations at a different frequency. With the advanced experiments and models, we will have the opportunity to take the full potential of this fast temperature tunable microfluidic device. These studies could help to investigate which hypothesis, i.e., cross-regulation by downstream genes or adaptive regulation from the upstream morphogen gradient, can account for the robust patterning in fly embryos under temperature perturbations.

## SUPPORTING MATERIALS

Supporting Materials include 8 supplementary figures, 3 supplementary tables, and 3 supplementary movies.

## AUTHOR CONTRIBUTION

Conceptualization: FL

Investigation: HZ YC

Methodology: HZ YC

Supervision: FL CL

Writing - original draft: FL HZ

## ACKNOWLEDGEMENTS

This work was supported by the National Natural Science Foundation of China [31670852, 11674010 and 11434001]. The *Drosophila* lab used in this project was supported by Peking-Tsinghua Center for Life Sciences.

